# A generalized adaptive harvesting model exhibits cusp bifurcation, noise, and rate-associated tipping pathways

**DOI:** 10.1101/2022.12.01.518756

**Authors:** E.W. Tekwa, Victoria Junquera

## Abstract

The sustainability of renewable resource harvesting may be threatened by environmental and socioeconomic changes that induce tipping points. Here, we propose a synthetic harvesting model with a comprehensive set of socioecological factors that have not been explored together, including market price and stock value, effort and processing costs, labour and natural capital elasticities, societal risk aversion, maximum sustainable yield (*MSY*), and population growth shape. We solve for harvest rate and stock biomass solutions by applying a timescale-separation between fast ecological dynamics and slow institutional adaptation that responds myopically to short-term net profit. The result is a cusp bifurcation with two composite bifurcation parameters: 1. consumptive scarcity *λ*_*c*_ or the ratio of market price-to-processing cost divided by *MSY* (leading to a pitchfork), and 2. non-consumptive scarcity *λ*_*n*_ or the stock value minus a scaled effort cost (leading to saddle-nodes or folds). Together, consumptive and non-consumptive scarcities create a cusp catastrophe. We further identify four tipping phenomena: 1. process (harvest rate) noise-induced tipping; 2. exogenous (*λ*_*c*_) rate+process noise-induced tipping; 3. exogenous noise-induced reduction in tipping; and 4. exogenous cycle-induced reduction in tipping. Case 2 represents the first mechanistically motivated example of rate-associated tipping in socioecological systems, while cases 3 and 4 resemble noise-induced stability. We discuss the empirical relevance of catastrophe and tipping in natural resource management. Our work shows that human institutional behaviour coupled with changing socioecological conditions can cause counterintuitive sustainability and resilience outcomes.

## Introduction

The harvesting of renewable resources is a transdisciplinary topic spanning ecology, environment, and economics (Heal, 1998; Ostrom, 2009; C. W. Clark, 2010; S. Levin et al., 2013; Dasgupta, 2021). An archetypal model of harvesting involves logistic growth of a resource coupled to a mortality or harvest rate caused by human institutions (Schaefer, 1954). While many variations have arisen in ecology, economics, fishery, forestry, and agricultural sciences to highlight complex dynamics and viable policies (Vernon L. Smith, 1968; Dasgupta, 1982; Morey, 1986; Heal, 1998; C. W. Clark, 2010; Costello et al., 2016), a synthetic model that captures various aspects of growth, benefit, cost, and human behaviour remains to be formalized. For example, most fishery models focus on harvesting cost due to effort (Gordon, 1954; C. W. Clark, 2010), but recent empirical syntheses (Lam et al., 2011) and models (Dasgupta et al., 2019; Tekwa et al., 2019) suggest that processing cost can be a major driver of socioecological dynamics. As well, bioeconomic models of harvesting have treated elasticities of labour and natural capital (the stock) using various functions that have so far been treated separately (Morey, 1986; Dasgupta et al., 2019). Finally, bifurcation analyses stemming from different models point to the potential for many different parameters (such as commodity prices, factor elasticities, or natural resource growth dynamics) and parameter dynamics (such as rapid rate of change or noise) to cause tipping point phenomena (Skiba, 1978; Crépin, 2007; Horan et al., 2011; Ashwin et al., 2012; Lade et al., 2013). These factors complicate management for marine (P. S. Levin & Möllmann, 2015; Selkoe et al., 2015; Sguotti et al., 2019), agro-pastoral (Tittonell, 2014; Lee et al., 2015), and water sustainability (Dunn et al., 2017). However, it remains unclear what are the generic parameter groups that cause similar dynamics.

Here we produce a synthetic model that captures ecological dynamics and includes a comprehensive set of benefits and costs of harvesting. We then reduce the parameters to interchangeable and identifiable variables and ask what the dynamic outcomes are when institutions adapt slowly (relative to ecology) and myopically to maximize economic yield (Tekwa et al., 2019). This procedure allows us to identify that harvesting generically exhibits a cusp bifurcation (Thom, 1969) with two scarcity bifurcation variables that capture different aspects of ecological and economic conditions. Finally, we explore the tipping implications of this model under exogenous changes in price, cost, and ecology, such as through trade networks and climate change (Ashwin et al., 2012). We find that bifurcation, exogenous and process noises, rates of change in exogenous variables, and exogenous cycles contribute to different types of tipping phenomena. In particular, we find that fast cycles or high exogenous noise reduce the chance of process noise-induced tipping, a “hula loop” effect that differs from previous tipping mechanisms involving cycles (Alkhayuon et al., 2021). The work provides a platform with which to explain and inform how changing socioecological conditions and institutions create a complex adaptive landscape for resource sustainability.

## Methods

### Model Definition

The socioecological system consists of two ordinary differential equations. Symbols for all state variables and parameters are defined in Table 1. The first equation describes the ecological dynamics with stock size *S* according to a Schaefer harvest function (Schaefer, 1954) with Pella-Tomlinson population dynamics (Pella & Tomlinson, 1969; Thorson et al., 2012; Costello et al., 2016):

**Table 1.** Symbol definitions. Empty entries under unit are dimensionless. For ranges, [ ] means including the value, while () means up to but not including the value.

| Symbol | Description | Unit | Range |
| --- | --- | --- | --- |
| $A$ | $\lambda_c$-cycle amplitude | | [0,∞) |
| $E$ | effort | | [0,∞) |
| $F$ | harvest rate | yr<sup>-1</sup> | [0,∞) |
| $FS$ | harvest volume or consumption | kg/yr | [0,∞) |
| $R$ | rate of directional change in $\lambda_c$ | yr<sup>-1</sup> | (-∞,∞) |
| $S$ | stock size | kg | [0,∞) |
| $MSY$ | maximum sustainable yield | kg/yr | [0,∞) |
| $a$ | intraspecific resource competition | kg<sup>-1</sup>yr<sup>-1</sup> | (0,∞) |
| $dW_t$ | Wiener process (stochasticity) | yr<sup>-1</sup> | [0,∞) |
| $d_Y$ | pitchfork imperfection term | \$<sup>3</sup>kg<sup>-3</sup>yr<sup>-3</sup> | (-∞,∞) |
| $f$ | $\lambda_c$-cycle frequency | yr<sup>-1</sup> | [0,∞) |
| $q$ | catchability | yr<sup>-1</sup> | [0,∞) |
| $r$ | intrinsic growth rate | yr<sup>-1</sup> | [0,∞) |
| $u$ | utility | \$/yr | (-∞,∞) |
| $\tilde{u}$ | simplified utility | \$/yr | (-∞,∞) |
| $w$ | rate of price change | yr/kg | [0,∞) |
| $u_{ES} = V_{ES}q^{-1/\alpha}$ | reduction in cost of effort by stock size | \$yr<sup>1/\alpha</sup>-<br>1/kg | [0,∞) |
| $u_E = -C_E q^{-1/\alpha}$ | cost of effort | \$yr<sup>1/\alpha-1</sup> | (-∞,0] |
| $u_S = V_S$ | stock value from ecosystem service | \$kg<sup>-1</sup>yr<sup>-1</sup> | (-∞,∞) |
| $u_{FS} = -C_{FS}$ | cost of processing harvested volume | \$/kg | (-∞,0] |
| $u_{\beta FS} = V_{\beta FS}$ | market price multiplier | \$/yr | [0,∞) |
| $\alpha$ | effort elasticity; $\alpha(1-\beta)$ is the labour elasticity | | (-∞,∞) |
| $\beta$ | society's risk aversion; $1-\beta$ is the natural capital elasticity | | (-∞,∞) |
| $\epsilon$ | adaptation rate | | →0 |
| $\rho$ | Pella-Tomlinson growth shape parameter | | (0,∞) |
| $\lambda$ | scarcity or shadow price | \$kg<sup>-1</sup>yr<sup>-1</sup> | [-∞,∞] |
| $\lambda_c = \frac{(wu_{\beta FS}/-u_{FS})}{wMSY}$ | (specific) consumptive scarcity | | [0,∞] |
| $\lambda_{c\beta} = \frac{(wu_{\beta FS}/-u_{FS})^{1/\beta}}{wMSY}$ | general consumptive scarcity | | [0,∞] |
| $\lambda_n = u_S - au_E$ | non-consumptive scarcity | \$kg<sup>-1</sup>yr<sup>-1</sup> | (-∞,∞) |
| $\sigma_F$ | standard deviation of process noise (in $F$) | yr<sup>-1</sup> | [0,∞) |
| $\sigma_{\lambda_c}$ | standard deviation of exogenous noise (in $\lambda_c$) | | [0,∞) |

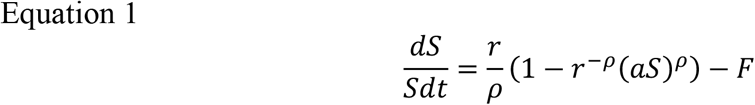

Per-capita growth rate is a function of intrinsic growth rate *r*, the Pella-Tomlinson growth shape parameter *ρ*, the intraspecific competition between harvested organisms *a*, and the harvest rate *F. FS* is the consumption rate. When *ρ=*1, we obtain logistic growth. When *ρ*<1, maximum sustainable yield (*MSY*) occurs closer to zero stock; and when *ρ*>1, *MSY* occurs closer to carrying capacity. Empirical studies show *ρ* is close to or slightly less than one (Thorson et al., 2012). The equilibrium biomass *S*^*\**^ given *F* is:

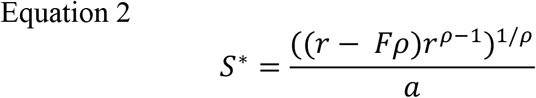

*MSY* is by definition *F*^*MSY*^*S*^*MSY*^, which occurs when *∂*(*dS/dt*|^*F=0*^)/*∂S*=0. Solving for *S* gives the following parameters:

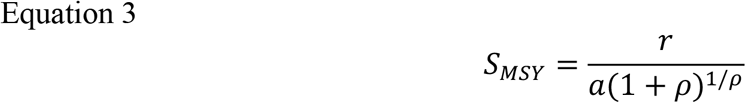

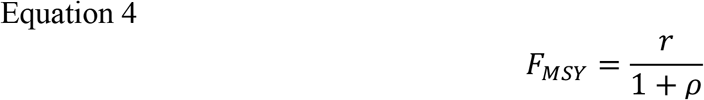

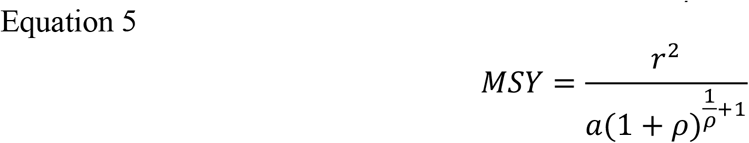

For *ρ=*1, we recover the standard solutions for logistic growth: *S*_*MSY*_=*r/*(2*a*), *F*_*MSY*_=*r*/2, and *MSY*=*r*^*2*^/(4*a*).

The second dynamic equation is derived from the following definition of utility *u*, which we first write in terms of various commonly measured benefits and costs:

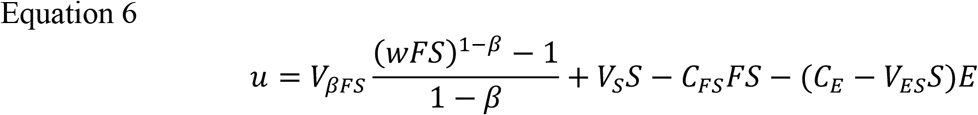

The first term on the r.h.s. of Equation 6 is an isoelastic function of consumption *FS* representing a non-linear form of utility from harvesting, which is the direct benefit from harvest (Dasgupta et al., 2019). *w* controls the rate of price change, and *β* controls the natural capital elasticity (see below for derivation) and can also be understood as a measure of societal risk aversion, since a higher *β* means higher value placed on low instead of high harvest volume (Cobb & Douglas, 1928; Morey, 1986). *V*_*βFS*_ controls the magnitude of marginal benefits and can be thought of as the exogenous market price multiplier.

The second source of benefit is from the standing stock, which is biomass times the intrinsic value or stock price *V*_*S*_ (second term in Equation 6). This stock price is commonly an externality not included in market valuation of the resource when benefits from standing biomass is not exclusive to harvesters. In this case stock price can only be valued cooperatively by a society that considers ecosystem services derived from natural capital, including the species’ contributions to ecosystem health, tourism, carbon sequestration, cultural significance, etc. (Heal, 1998; Dasgupta, 2021).

The third term in Equation 6 represents processing cost, which is a constant marginal cost *C*_*FS*_ times catch volume (Lam et al., 2011; Dasgupta et al., 2019; Tekwa et al., 2019). A commonly considered additional cost is the cost of effort, which is a constant marginal cost *C*_*E*_ times effort *E*, measured in labour years spent harvesting divided by a standardizing number of years – resulting in a dimensionless unit (in fourth term of Equation 6). The net marginal cost of effort may diminish with stock size since it may be easier to locate abundant organisms within a labour year (Dasgupta, 1982; Ling & Milner-Gulland, 2006). A simple expression of of this diminishing cost is a constant marginal benefit *V*_*ES*_*S* proportional to *E* (in fourth term of Equation 6).

We simplify Equation 6 by combining the benefit and cost parameters into generic utility parameters. First, we note that *F*=*E*^*α*^*q* where *α* is the effort elasticity and *q* is catchability via classical definitions (Morey, 1986). Given that *F* can now be written as a function of *E*, we can rewrite the isoelastic function as:

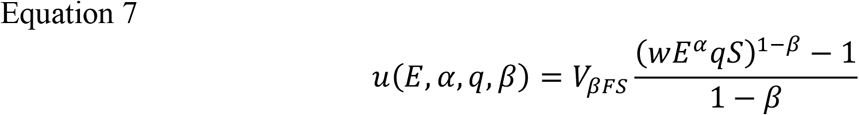

In this form, *α*(1-*β*) is the labour elasticity and 1-*β* is the natural capital elasticity in a classic Cobb-Douglas production function (Cobb & Douglas, 1928). When *β* is 0, there are no diminishing returns to consumption or harvest volume (i.e., price is proportional to harvest volume). When *β*<0, marginal returns increase with harvest volume. Equivalently, *β*<0 also describes a diminishing marginal cost with harvest volume – or economies of scale. Conversely, for *β*>0, marginal returns decrease with harvest volume, indicating diseconomies of scale or diminishing returns. The special case of *β=*1 results in a natural log function for diminishing returns.

The cost of effort (fourth term in Equation 6) can be written as (*C*_*E*_-*V*_*ES*_*S*)(*F/q*)^1/*α*^. This together with Equation 7 allows us to write Equation 6 only with *S* and *F* (eliminating *E*). Thus, we introduce the generic marginal utility components *u*_*βFS*_ (=*V*_*βFS*_), *u*_*S*_ (=*V*_*S*_), *u*_*FS*_ (=-*C*_*FS*_), *u*_*E*_ (=-*C*_*E*_*q*^-1/*α*^), and *u*_*ES*_ (=*V*_*ES*_*q*^-1/*α*^). In addition, we take out constant terms that do not affect the change in utility, which is what determines institutional dynamics as described later. The simplified utility function *ũ* is:

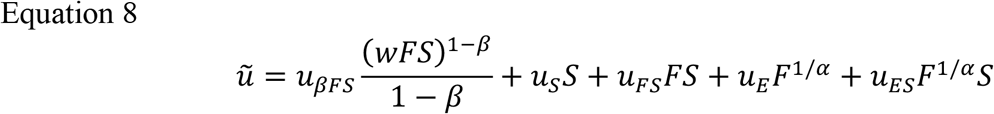

### Bifurcation Identification

We now assume that the institution adapts its harvest rate *F* incrementally through a myopic utility maximization (local gradient climb) process akin to evolution by natural selection (Dieckmann & Law, 1996; Tekwa et al., 2019). In this case, the change in *F* over time is proportional to the change in *u* in response to a change in *F* plus a stochastic component in *F*:

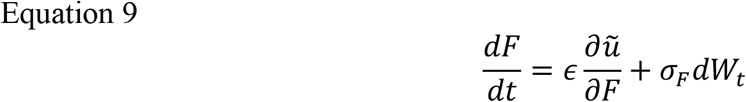

*ϵ* is the adaptation rate, *σ*_*F*_ is the standard deviation of noise in *F*, and *W*_*t*_ denotes a Wiener process – a one-dimensional Brownian motion or random walk with random step size – in a stochastic differential equation (Kloeden et al., 1997). In subsequent stochastic simulations of Equation 9 we will use the Euler-Maruyama method (Maruyama, 1955) where 𝒩(0, *Δt*) is a random number drawn from the normal distribution with mean zero and variance *Δt* (time step: 0.01 in simulations):

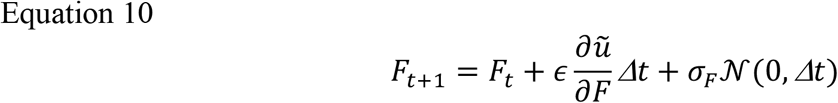

We then assume that institutions adapt slowly relative to ecological dynamics, i.e., ϵ→0, denoting a timescale separation. In this case, the ecological dynamics described in Equation 1 are always at equilibrium (*S*^*^) from the institutional perspective. Thus, we can solve for *S* in Equation 1, substitute the solution into Equation 9, and attain a one-dimensional system. The deterministic component of change in harvest rate can be rewritten as a polynomial in *F*, which reveals the normal form that identifies the bifurcation type (Supplementary Material: Appendix A).

We find that when *u*_*E*_, *u*_*S*_, and *u*_*ES*_ are zero, a pitchfork bifurcation occurs; else an imperfect pitchfork occurs – also known as a cusp bifurcation or catastrophe (Thom, 1969). This cusp bifurcation does not generally have closed-form solutions but can be solved numerically. The pitchfork bifurcation can be solved explicitly for *α=ρ=*1 and has the following set of at most three equilibria:

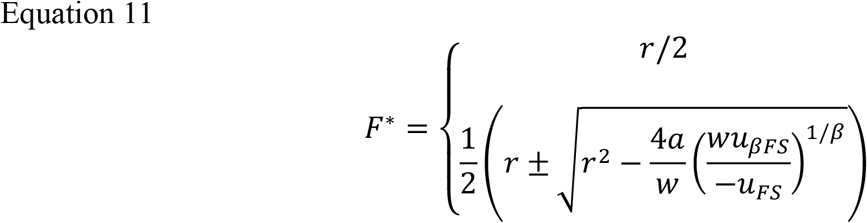

We solve for the stock biomass *S*/S*_*MSY*_ by substituting Equation 11 into Equation 1. The solutions in *F* and *S* are symmetric (with the 1+√ solution in *F* corresponding to the 1-√ solution in *S*). We then simplify the equilibria expressions using Equation 3 to Equation 5:

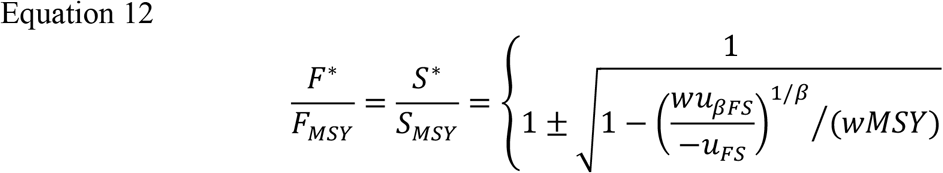

The stability of each solution *F*^*\**^ is indicated by *∂*^2^*u/∂F*^*2*^|_*F\**_<0. The upper stable biomass solution (or lower harvest rate solution) coincides with what is expected from optimization with a zero discount rate under the same socioecological conditions (Dasgupta et al., 2019; Tekwa et al., 2019).

We identify 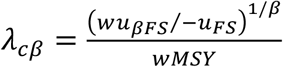 as the first of two general socioecological bifurcation parameters in the cusp bifurcation when *α=ρ=*1, since the number of real solutions in Equation 12 depends on whether *λ*_*cβ*_<1(three solutions) or *λ*_*cβ*_≥1 or is imaginary (one solution: *F*^*\**^*/F*_*MSY*_=1). The term *u*_*βFS*_/-*u*_*FS*_ can be understood as a ratio of market price-to-processing cost. *λ*_*cβ*_<1 indicates an economic constraint to harvest decision-making, since the condition implies the harvest is not valuable enough to maximize yield: at *MSY*, the resource has a shadow price or social scarcity value of zero (Dasgupta et al., 2019). In this case, the maximum economic yield (*MEY*) is less than the *MSY*. Conversely, when *λ*_*cβ*_≥1, the economic constraint to harvesting is released, with a positive scarcity value prompting the institution to adapt harvest rates based on ecological limitations. It can be shown that *λ*_*cβ*_ is related to a component of the shadow price that depends on consumption or harvest at *MSY*, thus we call *λ*_*cβ*_ the general “consumptive scarcity” (SM: Appendix B). A high consumptive scarcity *λ*_*cβ*_ means that price (*u*_*βFS*_) is high, processing cost is low (-*u*_*FS*_), maximum sustainable yield is low, and the rate of diminishing returns is low (*w*) when there are diminishing returns (*β*>0). When there are economies of scale (*β*<0), the above interpretation for high consumptive scarcity holds, except that now price (*u*_*βFS*_) is low and processing cost is high (-*u*_*FS*_). We choose to present the pitchfork results using a more specific consumptive scarcity parameter that is easier to grasp in terms of price and cost: 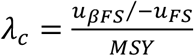 which leaves out the 1/*β* exponent in the numerator to be explored numerically for better intuition. Note *λ*_*c*_^*-*1^ was previously identified as a bifurcation parameter (Tekwa et al., 2019); here the parameter is inversed for improved comprehension.

*u*_*E*_, *u*_*S*_, and *u*_*ES*_ control the second dimension of the cusp bifurcation. The long-form expression of the resulting bifurcation parameter (*d*_*Y*_) when *α*=*β=ρ=*1 is derived in SM: Appendix A. A component of this bifurcation parameter is what we term the “non-consumptive scarcity”: *λ*_*n*_= *u*_*S*_ – *au*_*E*_ (SM: Appendix B). A high *λ*_*n*_ means the stock has high social value (*u*_*S*_) and is costly to harvest (-*u*_*E*_).

Matlab code to produce all simulation data and figures is available on https://github.com/EWTekwa/Tipping.

## Results

### Bifurcations

Here we show that the harvest model produces a cusp bifurcation or catastrophe under the assumption of slow and myopic institutional adaptation of harvest rate *F* based on the present utility gradient. In Figure 1, we focus on how the first bifurcation parameter, consumptive Scarcity 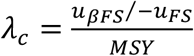, affects the harvested species biomass when other parameters are varied through eight combinations.

**Figure 1.**
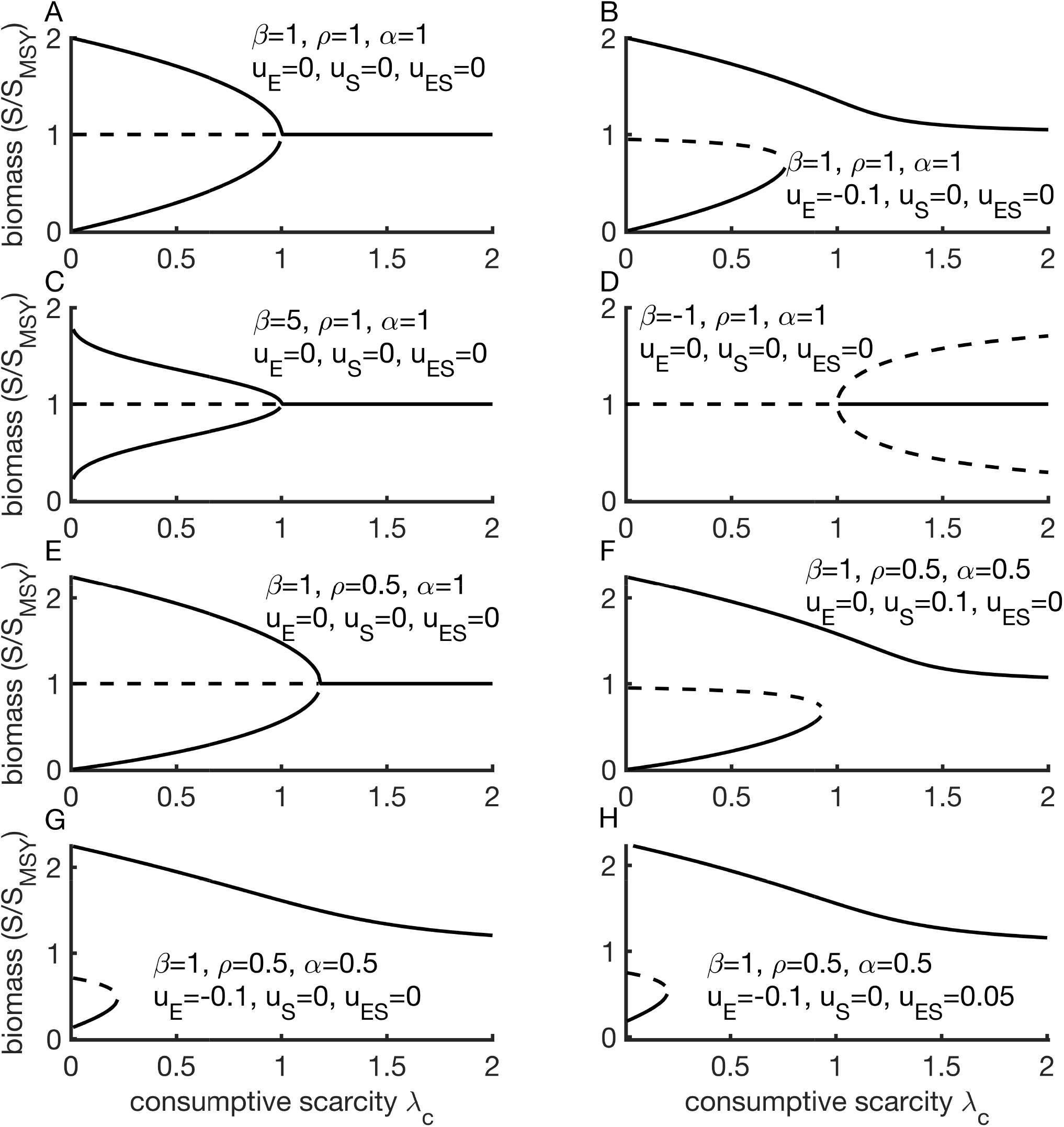
Numerical solutions for consumptive scarcity bifurcation. *λ*_*c*_ is a consumptive scarcity parameter (market price-to-processing cost ratio, *u*_*βFS*_*/-u*_*FS*_, divided by maximum sustainable yield, *MSY*). **(A-H)** *λ*_*c*_ bifurcations evaluated at various values for other parameters. Solid curves indicate stable equilibria, and dotted curves indicate unstable equilibria.

We see that throughout the parameter space explored, a pitchfork bifurcation occurs in *λ*_*c*_. An incremental change in the bifurcation variable *λ*_*c*_ produces a bifurcation-induced tipping or discontinuous change in biomass. Non-zero values in *u*_*E*_, *u*_*S*_, and *u*_*ES*_ produce a perturbed pitchfork by breaking the symmetry between the upper and lower solutions – or in a two-variable bifurcation perspective, a cusp bifurcation or catastrophe (Thom, 1969) with *λ*_*c*_ and one of *u*_*E*_, *u*_*S*_, and *u*_*ES*_ being the bifurcation variables (Figure 1A, B, F-H). This cusp will be clarified shortly. Growth characteristics (where *MSY* occurs relative to maximum biomass), controlled by *ρ*, also breaks symmetry (Figure 1E vs. 1A). Society’s risk aversion *β*, alternatively interpreted as the natural capital elasticity (1-*β*) as we showed in the Methods (Equation 7), controls whether the pitchfork bifurcation is supercritical (*β*>0) or subcritical (*β*<0) (Figure 1C-D).

Bifurcation-induced tipping can also occur along a second bifurcation dimension: the non-consumptive scarcity *λ*_*n*_= *u*_*S*_ – *au*_*E*_. Figure 2 shows that as *λ*_*n*_ changes between negative and positive, two saddle-node (i.e., fold) bifurcations meet. Such a sign change can come from sign change in either *u*_*S*_ or *u*_*E*_ when the other parameter is small. It is difficult to imagine *u*_*E*_ changing sign, since that would imply that harvest effort changes between being a cost and being a benefit (although bistability still exists for *u*_*E*_>0). However, *u*_*S*_ can reasonably change sign when a species changes from providing ecosystem services to becoming an ecosystem disservice. For example, a traditionally harvested species that is spatially shifting may become disruptive (decreasing *u*_*S*_) even when it retains a positive market value if consumed as captured by a positive *u*_*βFS*_. Conversely, an initially invasive species may in the long run provide positive ecosystem services (increasing *u*_*S*_). The entire cusp bifurcation across the two dimensions of *λ*_*c*_ and *λ*_*n*_ is shown in Figure 3.

**Figure 2.**
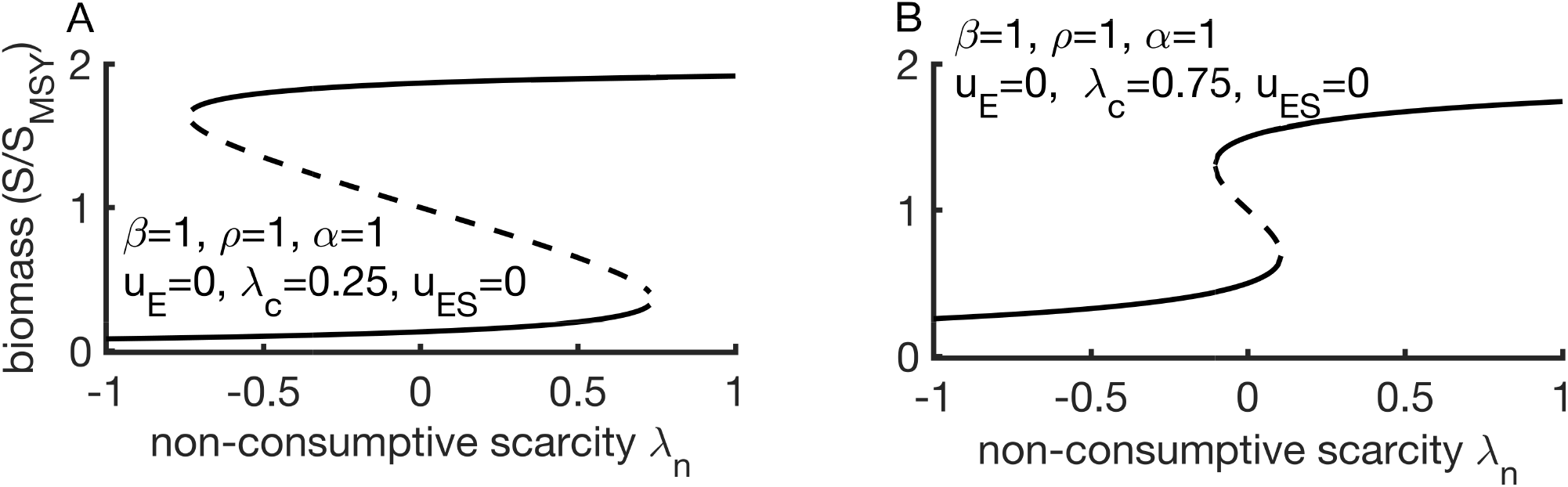
Numerical solutions for non-consumptive scarcity bifurcation. The non-consumptive scarcity parameter *λ*_*n*_ is intrinsic stock value *u*_*S*_ minus intraspecific competition *a* times cost of effort *u*_*E*_. **(A-B)** *λ*_*n*_ bifurcations evaluated at various values for other parameters. Solid curves indicate stable equilibria, and dotted curves indicate unstable equilibria.

**Figure 3.**
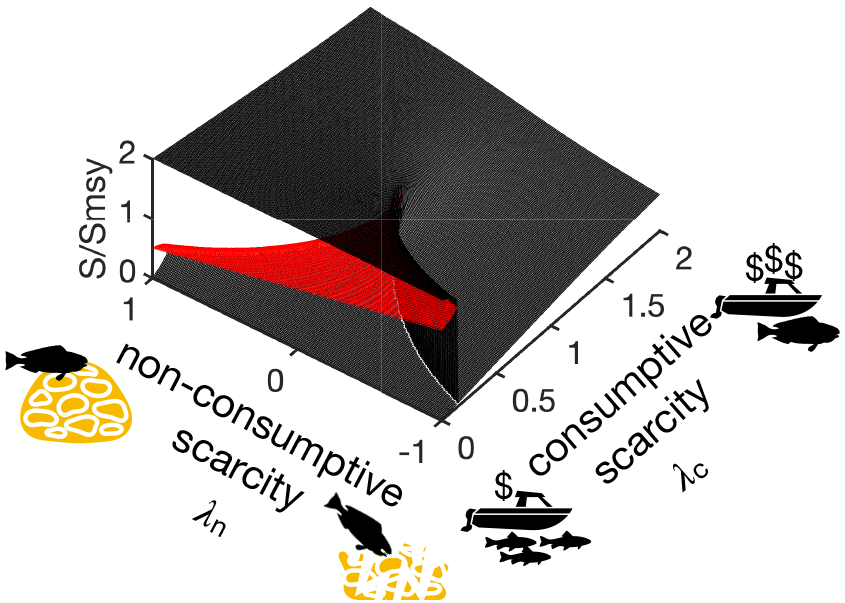
Numerical solutions for the cusp bifurcation. For *ρ=α=β=*1, relative biomass (*S/S*_*MSY*_) is determined by the two scarcity parameters *λ*_*c*_ (consumptive) and *λ*_*n*_ (non-consumptive). Black surface indicates stable equilibria, and red surface indicates unstable equilibria.

### Tipping Phenomena

We now explore different types of tipping phenomena across a range of *λ*_*c*_ values assuming a fixed negative *u*_*E*_ (Figure 4). In particular, we examine various scenarios of noise and rate-induced transitions. For each scenario, we simulate 100 stochastic trajectories with initial *F* being the higher stable biomass.

**Figure 4.**
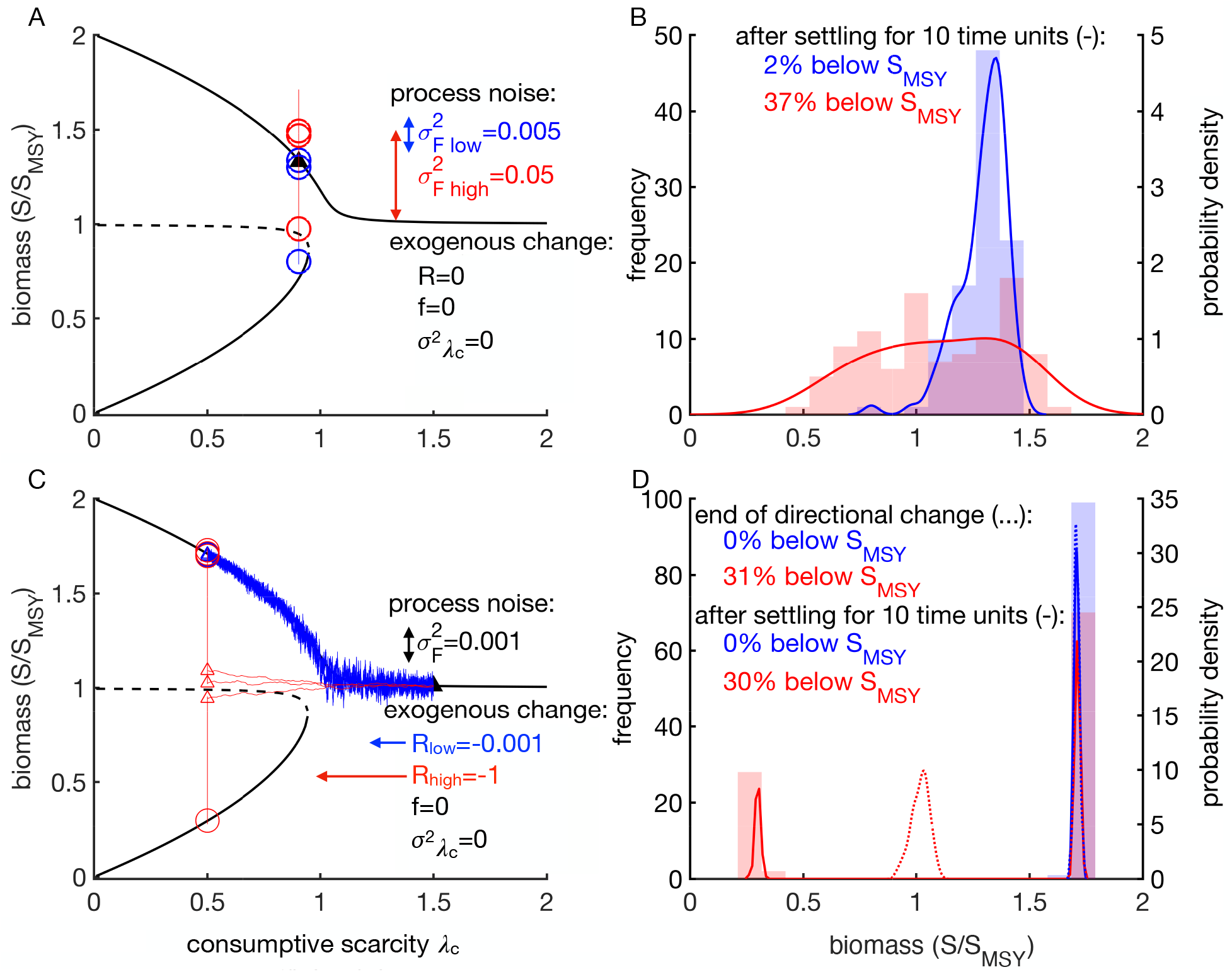
Process noise and rate-induced tipping. **(A & C)** Three example trajectories are illustrated for each of low noise (blue) and high noise (red) scenarios, with black triangle indicating the initial state and open circles indicating final states. Additionally for **C**, small open triangles indicate states at the end of directional exogenous changes. **(B & D)** Intermediate (dotted) and final (solid) biomass distributions for the low noise and high noise scenarios over 100 replicates each.

In addition to bifurcation-induced tipping (from a very slow change in *λ*_*c*_ as discussed in the previous section), we hypothesize that process noise can induce tipping because of stochasticity in harvest rate *F* (Equation 9). We further hypothesize that tipping can also be induced by fluctuation in the rate and direction of change of the exogenous parameter *λ*_*c*_. The change in the bifurcation variable (*dλ*_*c*_/*dt*) can be expressed as a combination of a rate of directional (or mean) change *R*(*t*), which may only be non-zero for some time *t*, a cyclical change with amplitude *A* and frequency *f*, and an exogenous noise *σ*_*λc*_ (which contrasts from process noise *σ*_*F*_).

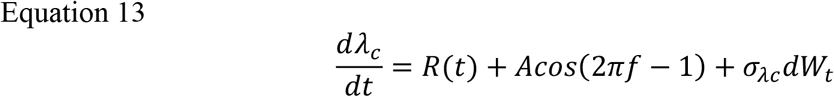

We first examine the effect of process noise (*σ*_*F*_ in Equation 9) for a constant *λ*_*c*_. A sufficiently large process noise (variance *σ*_*F*_^*2*^=0.05 compared to 0.005) causes a higher probability of tipping from an originally high biomass state to a low biomass state (Figure 5A-B red vs. blue). This is an example of a process noise-induced tipping.

**Figure 5.**
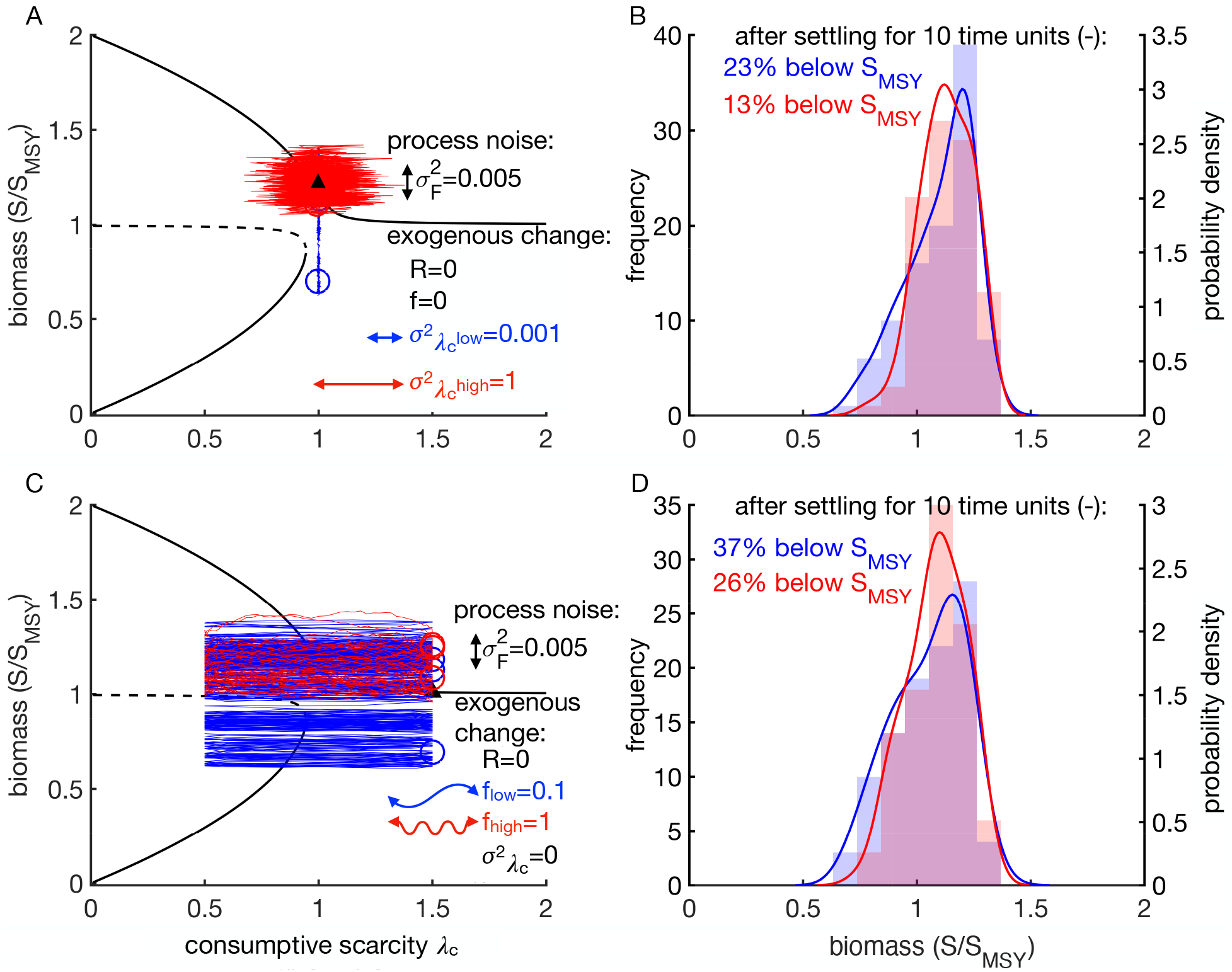
Exogenous noise and cycle-induced tipping. **(A & C)** Three example trajectories are illustrated for each of low and high fluctuation scenarios (blue vs. red), with black triangle indicating the initial state and open circles indicating final states. **(B & D)** Final biomass distributions for the low and high scenarios over 100 replicates each.

We next examine the effect of rates of change of the exogenous variable (*R*≠0). We first assume |*R*|*<<dS/dt*. That is, ecological dynamics, occurring at the fast timescale, equilibrate to the current harvest rate *F* and parameter value *λ*_*c*_ before these change. Additionally, assume a small process noise (*σ*_*F*_^*2*^=0.001). Then, a relatively high rate of decrease in *λ*_*c*_ from the single-solution regime at high values of *λ*_*c*_ (*R*=-1 vs. -0.001) followed by a period absent of directional change (*R*=0) results in a higher chance of tipping to a lower biomass (Figure 5C-D red vs. blue). The contrast between slow and fast changes demonstrates a rate+process noise-induced tipping in the socio-ecological system of harvesting. Note that here process noise is a necessary component for tipping because the deterministic upper biomass branch never dips below the lower biomass branch.

We now explore the role of exogenous noise *σ*_*λc*_ in tipping. Analogous to the previous case, exogenous noise alone cannot produce tipping in this system (not shown). However, a high level of exogenous noise (*σ*_*λc*_=1 in Equation 13) along with a sufficiently large process noise (*σ*_*F*_*=*0.005 in Equation 9) *decreases* the chance of tipping to a lower biomass relative to a small *σ*_*λc*_ (=0.001, Figure 5A-B red vs. blue). Under high exogenous noise, the institution has insufficient time to adapt to the current condition and instead sees the average condition (in this example *λ*_*c*_=1), which has a stable biomass solution above the unstable *S*_*MSY*_ solution. On the other hand, low exogenous noise gives the institution more time to adapt to the lower biomass equilibrium when process noise takes it to the lower basin of attraction. Subsequently, there is an elevated chance of remaining in the lower basin despite occasional escapes to *λ*_*c*_ conditions implicating higher biomass. High exogenous noise in this model may be considered a type of noise-induced stability (D’Odorico et al., 2005).

A similar phenomenon to noise-induced stability is also seen when the exogenous change is cyclical and process noise is present (Figure 5C-D). Again, the effect is moderate but a high frequency cycle reduces the chance of tipping to a lower biomass compared to a low frequency cycle (*f*=1 vs. 0.1, red vs. blue).

## Discussion

We incorporated into one synthetic harvesting model the salient ecological and economic parameters that have previously been studied in isolation. We discovered two composite bifurcation parameters. The first bifurcation group is the consumptive scarcity *λ*_*c*_*=(u*_*βFS*_/-*u*_*FS*_)/*MSY*, which is the consumptive price to cost ratio divided by maximum sustainable yield. This bifurcation parameter creates a pitchfork bifurcation in expected biomass due to an institution slowly adapting its harvest rate in a utility gradient climb. The second bifurcation parameter is the non-consumptive scarcity *λ*_*n*_, which is *u*_*S*_ (intrinsic stock value) plus *a* (intraspecific competition) times -*u*_*E*_ (cost per harvesting effort). This bifurcation parameter creates two joined saddle-nodes or folds in expected biomass. When these two bifurcation groups are combined, a cusp bifurcation or catastrophe is obtained. All other parameters stretch the shape of this basic 2-dimensional cusp biomass manifold. In particular, the species’ growth shape determines the parameter space in which bistable harvest rate occurs (such space increases when *ρ<*1, which is when *MSY* occurs at a stock lower than half of carrying capacity). On the other hand, the sign of societal risk aversion *β* (with 1-*β* being the natural capital elasticity) determines whether harvest experiences diminishing returns (*β>*0) or economies of scales (*β<*0), the latter flipping the cusp manifold stability. Effort elasticity *α* (or labour elasticity *α*(1-*β*)) had minor quantitative effects but we did not fully explore its effects; it is possible that some values can lead to additional equilibria.

The cusp catastrophe was traditionally used as a generic conceptual or mental model that has somewhat fallen out of fashion in biology and socioeconomics since its inception because of its frequent misapplications, often without mechanistic justifications or empirical evidence (Thom, 1977; Zahler & Sussmannt, 1977). However, the mathematics of cusp catastrophe can afford insights to complex dynamics when the bifurcation variables are mechanistically justified and make empirically testable predictions, as it has in physics. For example, some aspects of the visualization of gravitational lensing by black holes through cusp bifurcation, such as that seen in Interstellar (James et al., 2015), has since been supported by direct observations (Wilkins et al., 2021). Recent empirical applications of the cusp catastrophe can also be found in economics (Barunik & Vosvrda, 2009; Diks & Wang, 2016) and ecology (Sguotti et al., 2019) based on tailored statistical methods (Chow et al., 2015; Grasman et al., 2009), although mechanistic principles underlying these studies remain lacking.

Here we built a comprehensive harvesting model based on established principles in ecology and natural resource management. The emergent cusp bifurcation was unexpected and is worthy of closer investigation. One dimension of this cusp bifurcation – the effect of consumptive scarcity *λ*_*c*_ on harvesting decisions – has already been empirically supported by the bistable behaviour of global commercial marine fisheries (Tekwa et al., 2019). The other dimension, non-consumptive scarcity *λ*_*n*_, includes the cost of effort, which is the dominant cost considered and demonstrated in classical bioeconomic models that often do not consider *λ*_*c*_ (Heal, 1998; C. W. Clark, 2010). A catalogue of bifurcation types has been documented in resource harvest models (Ludwig et al., 1997; Crépin, 2003; Bauch et al., 2016), but to our knowledge our model is the first to find a cusp bifurcation in socioecological systems.

Our work explores the roles of process and exogenous noises, rate of change, and cyclic variability in socioecological tipping. Our model shows that process noise (stochasticity in harvest rate) can induce tipping. We further show that tipping can be induced by a combination of rapid decrease in consumptive scarcity and process noise. Such a situation might emerge, for example, when market prices decreases due to global trade, processing cost increases due to supply chain disruption, or *MSY* increases due to global warming and increased metabolic rate (by no mean true for all species). The latter is particularly surprising because increased growth rate, which by itself may be expected to improve sustainability, is here a cause of overharvesting (if such an increase is attained quickly).

Counterintuitively, our results also reveal that process noise coupled with high exogenous noise or rapid cycle in consumptive scarcity such as through price, cost, and *MSY* volatilities, can decrease the chance of tipping. These phenomena are related to noise-induced stability, which has been shown in ecosystems (D’Odorico et al., 2005) and physical systems (Mantegna & Spagnolo, 1996), but to our knowledge has not been shown in socioecological systems. Moreover, the decomposition of consumptive scarcity variability into directional change, cyclical change, and noise connects socioecological modelling to a rich literature on economic volatility (Mackey, 1989; Hu & Øksendal, 2003; Zilberman et al., 2013; Karali & Power, 2013).

Exogenous socioecological changes are rampant due to developments in trade, technology, disaster, politics, and climate change. For example, international trade can increase market prices (*u*_*βFS*_) for some exporters and decrease prices of substitutable resources for importers (Elsler et al., 2022). Market prices may also change due to marketing campaigns on previously ignored species, such as the orange roughy in New Zealand (M. Clark, 1995), or on climate change-induced novel species, such as lionfish in the US (Nuñez et al., 2012). Processing cost (-*u*_*FS*_) may increase due to rises in labour wage among processors, transportation cost, fuel cost, or cold storage cost (Brennan, 1976; Banoub & Martin, 2020) because of wars and pandemics; these costs may also drop due to technological advances. Analogous changes to cost of effort (*u*_*E*_) would result from impacts on harvesters instead of processors. *MSY* may increase for some species and decrease for others due to climate change, for example through metabolic response (Bernhardt et al., 2018) and range shift-mediated community reshuffling (Tekwa et al., 2022). Stock value (*u*_*S*_) may change or even flip sign because of changing ecosystem services or disservices that a species provides. Spatially shifting otters on the west coast of North America may disrupt valued seafood supply and thus accrues a negative stock value for some communities, even as the species retains a positive market price from consumption and provides tourism (Burt et al., 2020). Land use changes in Laos may cause previously valued rodents and plants to become pests (Rasmussen et al., 2017). These changes can occur at different rates, amplitudes, and frequencies, can be encapsulated into changes in the scarcity parameters *λ*_*c*_ and *λ*_*n*_, and can move a socioecological system along all directions of a cusp catastrophe leading to various tipping phenomena. Therefore, our theory predicts the resilience of societies facing multidimensional changes today (Nyström et al., 2019).

Our model has several shortcomings. First, we assumed a timescale separation between slow institutional adaptation and exogenous changes relative to fast ecological dynamics. While many commercial and governmental management institutions likely satisfy this timescale assumption (Pershing et al., 2015; Tekwa et al., 2019), other more nimble institutional types like cooperatives and loose collectives of individual harvesters may act as fast as the ecological timescale, in which case the cusp bifurcation dynamics does not occur. If exogenous changes occur at a similar timescale as ecological dynamics, then the rate-associated tipping phenomena we presented may also be invalid. We expect fast institutions to result in open-access like behaviour (Tekwa et al., 2019), but this has not yet been fully addressed along both scarcity dimensions introduced in the current paper. Exploring variations in institutional versus ecological timescales remain a research frontier (Crépin, 2007; Fryxell et al., 2010; S. Levin et al., 2013). Future research may more fully explore timescale issues using computer simulations and mathematical timescale separation techniques.

The second caveat to our results is that they are limited to myopic gradient-climbing adaptive institutions, which is separate from the issue of how fast institutions adapt. The myopic perspective means institutions look backward in time to determine what is the best harvest strategy that yields an increasing utility, which does not include a forward-looking component typical in other bioeconomic models through the computation of discounted net present value. While there is evidence that institutions behave more like myopic adaptive agents than forward-looking optimization agents (Selten, 1990; Nelson & Winter, 2002; David, 2007; Tekwa et al., 2019), reality is likely represented by diverse institutions that span a spectrum and may compete for a common resource (Ostrom, 2005). Both theory and empirical applicability will be improved by considering whether the cusp bifurcation in harvest rate still holds when institutions adapt differently and are heterogeneous within one system.

The third caveat is that while our set of socioecological parameters is more comprehensive than most harvesting models, it does not explicitly include sunk costs, or one-time investments that cannot be reversed. Some examples include purchasing a specialized fleet or equipment to increase the maximum attainable harvest rate, or legislating quotas or species protection status (Sims & Finnoff, 2016). While sunk costs can be used to justify the slow institutional dynamic assumption, it may be possible to explicitly incorporate sunk costs into the utility calculus. But doing so would imply adopting a discounting framework (otherwise an institution may never invest in harvesting capability if the immediate return does not cover investment cost). Connecting the backward-looking adaptive perspective with the forward-looking optimization perspective will clarify how socioecological systems evolve.

Our discovery of the cusp bifurcation in harvesting systems unfolds socioecological dimensions that were previously unintegrated. Limitations in modelling institutional adaptation imply that other dynamic outcomes are likely. Nevertheless, our results illustrate that coupled social and ecological dynamics can create sustainability consequences that counter expectations obtained from considering factors in isolation.

## Acknowledgements

ET is supported by a Hakai Postdoctoral Fellowship through the Tula Foundation. VJ is supported by Princeton University’s High Meadows Environmental Institute, Dean for Research, Andlinger Center for Energy and the Environment, and the Office of the Provost. We thank Vitor Vasconcelos, Flavia Marquitti, Lisa McManus, and Teresa Ong for organizing the Banff International Research Station workshop on *Rate-induced transitions in networked systems* where this paper was conceptualized.

## Appendix A. Normal Form

Here we derive the normal form for the change in harvest rate *dF/dt*, which is proportional to *∂ũ/∂F* (Equation 9). We first compute *∂ũ/∂F* from *ũ* (Equation 8) and rearrange terms so that the r.h.s. is grouped by polynomial orders in *F*:

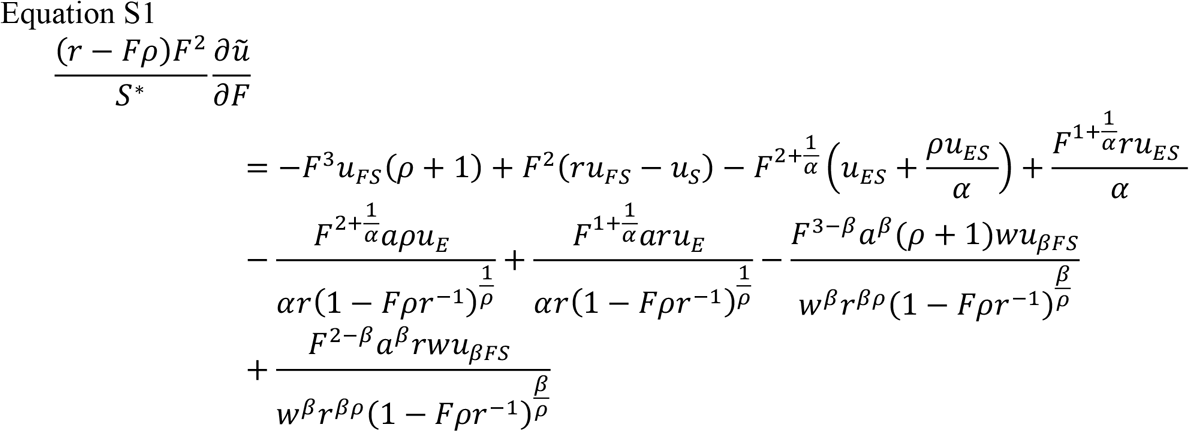

We see that there is always a cubic term in the state variable *F*, which implies three possible equilibria (solving for *F* in Equation S1). The presence of a cubic term plus a linear term (*F*^*3*^and *F*) implies a pitchfork bifurcation, while a cubic term plus a constant implies a cusp bifurcation (also known as an imperfect pitchfork) (Strogatz, 2015). Additional equilibria are possible when the exponents *α, β*, or *ρ* do not equal one. For the simple case of *α*=*β=ρ=*1, we can obtain a polynomial form with integer exponents in *F*:

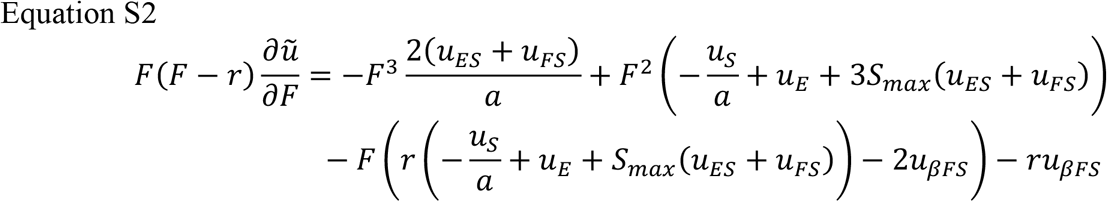

We now consider the case of *u*_*E*_=*u*_*S*_=*u*_*ES*_=0. Under these assumptions, Equation S2 takes the form:

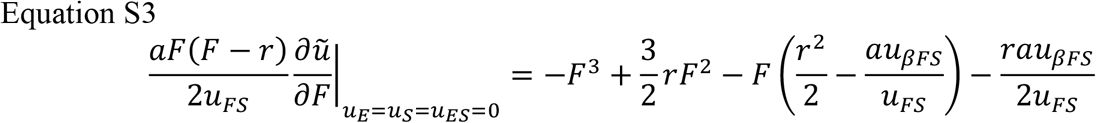

Any cubic equation can be reduced into a depressed cubic form through a change of variable (*F*=*Y-b/(3a)* where *a* and *b* are coefficients of *F*^*3*^ and *F*^*2*^). The depressed cubic form of Equation S3 has no *Y*^*2*^ or constant terms, which is the normal form of a pitchfork bifurcation (*dY/dt*=*Y*^*3*^*+b*_*Y*_*Y* where *b*_*Y*_ is a coefficient). Thus we can deduce that the bifurcation parameter controlling the pitchfork bifurcation includes *u*_*βFS*_ and *u*_*FS*_ but not *u*_*E*_, *u*_*S*_, and *u*_*ES*_.

In contrast, when the terms *u*_*E*_, *u*_*S*_, or *u*_*ES*_ are non-zero, Equation S2 is of the form *dY/dt*=*Y*^*3*^*+b*_*Y*_*Y*+*d*_*Y*_ (where *d*_*Y*_ is a non-zero constant) which is an imperfect pitchfork – also known as the normal form of a cusp bifurcation (Thom, 1969). *d*_*Y*_ is the imperfection term and a bifurcation parameter because it controls whether bistability occurs; in particular it creates a hysteresis loop where two fold bifurcations join:

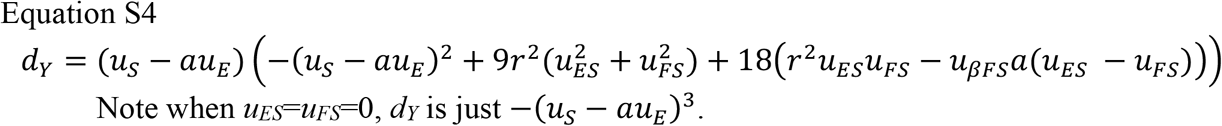

## Appendix B. Shadow Price

The shadow price or social scarcity value *λ* per consumption unit (*FS*) is defined as *∂ũ/∂FS*. In other word, *λ* is the marginal utility to society of consuming an additional amount given the current conditions (*F, S*), which is not the same as the market value of consumption. *λ* is obtained from taking the partial derivative of Equation 8 with respect to *FS*:

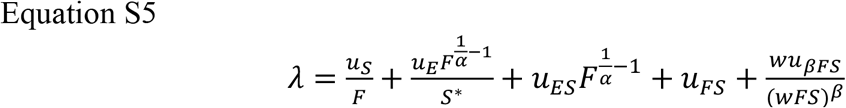

A transformation of shadow price associated with *u*_*FS*_ and *u*_*βFS*_ (i.e. when *u*_*S*_=*u*_*E*_=*u*_*ES*_=0) gives us a “consumptive scarcity” variable *λ*_*FS*_:

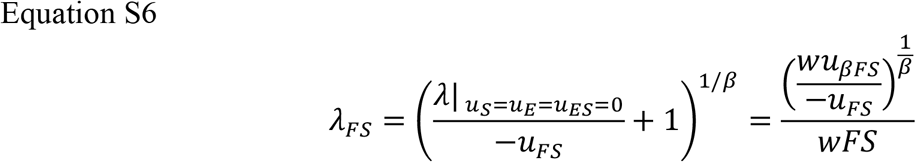

Note this is a monotonic (order-preserving) transformation only when *β*>0. This consumption scarcity variable computed at *FS=MSY*, hereby labelled *λ*_*cβ*_, turns out to be a bifurcation parameter as identified in the main text:

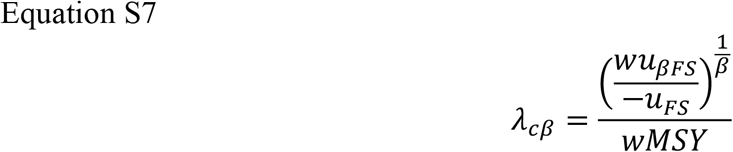

On the other hand, the shadow price associated with *u*_*S*_, *u*_*E*_, and *u*_*ES*_ gives us a “non-consumptive scarcity” variable:

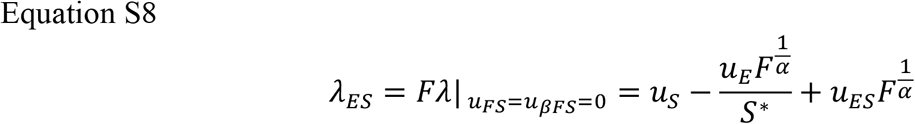

In particular, when *α*=1, *ρ*=1, *u*_*ES*_=0, and *F=F*_*MSY*_=*r* (following the above observation that *MSY* is the key consumption rate), the non-consumptive scarcity (let’s call *λ*_*c*_) is:

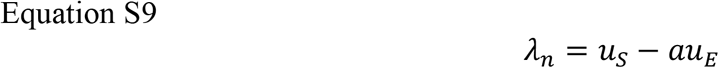

By inspection, *λ*_*n*_ is proportional to the bifurcation parameter *d*_*Y*_ (the second bifurcation dimension of the cusp, see Equation S4). Thus, while non-zero values *u*_*ES*_ and *u*_*FS*_ will modify how *λ*_*n*_ affects the *F* equilibria, *λ*_*n*_ alone can also be considered a bifurcation parameter. In the main text we use *λ*_*n*_ as the second bifurcation dimension in the cusp, since this form has a clear socioecological meaning. A high *λ*_*n*_ means the stock has high social value and is costly to harvest.

